# A method for the unbiased quantification of reassortment in segmented viruses

**DOI:** 10.1101/708982

**Authors:** Megan R. Hockman, Kara Phipps, Anice C. Lowen

**Affiliations:** Department of Microbiology and Immunology, Emory University School of Medicine, Atlanta, Georgia, United States of America

**Keywords:** Reassortment, reovirus, influenza virus, viral genetics

## Abstract

The diversification of segmented viruses via reassortment is important to understand due to the contributions of reassortment to viral evolution and emergence. Methods for the quantification of reassortment have been described, but are often cumbersome and best suited for the analysis of reassortment between highly divergent parental strains. While it is useful to understand the potential of divergent parents to reassort, outcomes of such heterologous reassortment are driven by differential selection acting on the progeny and are typically strain specific. To quantify reassortment, a system free of differential selection is needed. We have generated such a system for influenza A virus and for mammalian orthoreovirus by constructing well-matched parental viruses carrying small genetic tags. The method utilizes high-resolution melt technology for the identification of reassortant viruses. Ease of sample preparation and data analysis enables streamlined genotyping of a large number of virus clones. The method described here thereby allows quantification of the efficiency of unbiased reassortment and can be applied to diverse segmented viruses.

**Highlights:**

- Genetic tagging of viruses can be achieved without altering fitness
- High-resolution melt can detect single nucleotide differences in viruses
- Unbiased reassortment of influenza A virus and mammalian orthoreovirus can be quantified

## Introduction

Virus genome organization varies widely across different families. Segmented genomes, which are comprised of distinct RNA or DNA molecules, have been documented in eleven virus families to date. Segment numbers vary, as do replication strategies. Each genome segment encodes a different protein or proteins which are involved in establishing a productive infection. During infection, cells can be co-infected by multiple parent viruses. In segmented viruses, co-infection events have the potential to give rise to many different progeny genotypes, fueling evolution.

Viruses with segmented genomes undergo a type of genetic recombination termed reassortment. Reassortment occurs when two or more parent viruses co-infecting a single cell exchange whole gene segments, which are then packaged together to yield progeny viruses with novel genotypes. This mode of generating diversity is unique to segmented viruses, and contrasts with classical recombination, in which a chimeric product is formed from the breaking and rejoining of nucleic acid strands. Reassortment events most often yield attenuated progeny due to incompatibilities between nucleic acids and/or proteins derived from heterologous parents^1-5^. There is, however, the potential for the coupling of compatible, beneficial alleles and the subsequent emergence of novel pathogens, as was observed in the 2009 influenza A virus (IAV) pandemic^6, 7^.

To date, reassortment has been investigated in several viruses, including IAV and mammalian orthoreovirus, using a variety of methods. Measurement of reassortment is often complicated by fitness differences among progeny viruses or a lack of sensitive quantification methods. In the former case, difficulties arise due to selection. When reassortment yields progeny of variable fitness, those with the most beneficial segment combinations will be amplified more rapidly. Preferential propagation of parental genotypes results in under-estimation of the frequency of reassortment. Similarly, preferential amplification of certain reassortant viruses results in under-estimation of population diversity. To avoid these issues, it is preferable to quantify reassortment between viruses of more similar genotypes. This can be challenging, however, as most detection methods rely on parental viruses being significantly different genetically. Detection of reassortment in genetically similar viruses requires sensitive molecular technologies, which were not available until relatively recently^8^.

The earliest method to identify reassortants utilized polyacrylamide gel electrophoresis^9-11^. This method depends on all segments of each parental genome having different electrophoretic mobility. For this to be the case, each segment must differ significantly in length and/or sequence. The requirement for significant differences for detection necessitates that parent viruses are divergent which can impact the fitness of reassortant progeny and reduce observed rates of reassortment. Additionally, there is a practical limitation on the number of samples that can feasibly be analyzed using this approach.

As an alternative approach, temperature-sensitive (ts) mutants have been used to quantify reassortment^12, 13^. In this system, a single segment confers temperature-sensitivity in each parental virus and pairs of viruses used for co-infection carry ts mutations in differing segments. Temperature sensitivity is abrogated if these segments are exchanged for wild type segments from the opposite parent in a reassortment event. Culture of progeny viruses at the nonpermissive temperature results in selection of wild type reassortants. Titration of progeny viruses at the nonpermissive temperature therefore allows quantification of the frequency of exchange of the ts segments. This approach makes it possible to detect reassortment between identical parents (with the exception of the ts lesions), thus avoiding the confounding effects of protein or segment incompatibilities. However, it does not yield the frequency of reassortment between all virus segments but rather only the two ts segments^12, 14^.

PCR-based methods can also be used to differentiate parental segments that differ sufficiently to allow the design of specific primers^15-17^. Amplicons can be detected using gel electrophoresis with ethidium bromide staining, or by determining C_t_ values in quantitative PCR. These methods, however, may be limited when parental genomes are too similar to allow unique primer design for all segments.

Alternatively, whole or partial genome sequencing of clonal virus populations offers a very flexible approach to identify reassortants^18^. With sequencing as a read-out, the need for primer design does not constrain the parental viruses that can be examined in combination. Traditionally, the costs of sequencing approaches have prohibited their use on a larger scale. In recent years, however, the price of sequencing has decreased and the technology has improved.

To address the shortcomings of prior methodologies and enable robust quantification of reassortment, we present here two methodological innovations. First, to eliminate selection bias, we generated well matched pairs of parental viruses that differ only by one or a handful of synonymous mutations in each genome segment. These introduced synonymous changes act as genetic markers to indicate the parental origin of each segment. Second, to streamline the detection of reassortant viruses, we applied high resolution melt analysis, a post-PCR method originally developed to differentiate single nucleotide polymorphisms in eukaryotic genomes^8^. Because this method is sensitive enough to detect single nucleotide differences, it allows for the quantification of reassortment between highly similar parental viruses. We have applied these approaches to both IAV and mammalian orthoreovirus.

## Materials and Methods

### Cell lines and cell culture media

293T cells from the American Type Culture Collective (ATCC) were maintained in Dulbecco’s Modified Eagle Medium (DMEM; Gibco) supplemented with 10% fetal bovine serum (FBS; Atlanta Biologicals). Madin-Darby Canine Kidney (MDCK) cells (from Dr. Peter Palese, Icahn School of Medicine at Mount Sinai) were maintained in Minimal Essential Medium (MEM; Gibco) supplemented with 10% FBS and 100U/100 μg/mL penicillin/streptomycin (Corning). Baby hamster kidney cells stably expressing T7 RNA polymerase (BHK-T7) cells^19^ were maintained in DMEM supplemented with 5% FBS, 2 mM L-Glutamine (Corning), 100U/100 μg/mL penicillin/streptomycin and 1 mg/mL G418 (Gibco). Spinner-adapted L929 cells (gift from Bernardo Mainou) were grown in Joklik’s modified MEM supplemented with 5% FBS, 2 mM L-Glutamine, 100U/100 μg/mL penicillin/streptomycin and 0.25 mg/mL amphotericin B (Sigma), termed SMEM.

### Design of wild-type and variant viruses

To allow quantitative analysis of reassortment, we designed parental viruses that are i) highly homologous, so that reassortment does not give rise to genetic or protein incompatibilities, and ii) genetically distinct in all segments to allow tracking of genetic exchange. For both IAV and mammalian orthoreovius, a variant virus was generated from the wild-type strain using reverse genetics. The wild type virus strains used were influenza A/Panama/2007/99 (H3N2) (Pan/99) virus and Type 3 Dearing (T3D) mammalian orthoreovirus. To generate variant (var) viruses, silent mutations were added to the first 1kb of each wild-type virus coding sequence, as shown in Table 1. We have made multiple versions of the Pan/99var virus^20, 21^; the Pan/99var virus described here is Pan/99var15. The mutations made were A to G/G to A or C to T/T to C, which have the greatest impact on melting properties^8^. The melting properties of a short amplicon (65-110 base pairs) containing a mutation of this nature is typically altered sufficiently to allow robust detection by high resolution melt analysis^8, 22^. To avoid introducing attenuating mutations, sites of natural variation were targeted where sequence data was available. For IAV, isolates from the same lineage within a 10 year time frame were selected from the NCBI GenBank database, and 20-30 sequences were aligned. Sites within the first 1000 bp relative to the 3’ end of the vRNA that displayed high nucleotide diversity were targeted for introduction of variant mutations. Point mutations were introduced into plasmid-encoded viral cDNAs using QuikChange mutagenesis (Agilent) according to the manufacturer’s protocol.

**Table 1:** Point mutations used to generate variant viruses.

| T3D Mammalian Orthoreovirus |  | Pan/99 Influenza A Virus |  |
| --- | --- | --- | --- |
| Segment | Polymorphism | Segment | Polymorphism(s)* |
| L1 | C612T | PB2 | C354T C360T |
| L2 | C853T | PB1 | A540G |
| L3 | G481A | PA | A342G G333A |
| M1 | C919T | HA | T308C C311A C314T A646T<br>C467G T470A |
| M2 | A650G | NP | C537T T538A C539G |
| M3 | T702C | NA | C418G T421A A424C |
| S1 | G312A | M | G586A |
| S2 | A438G | NS | C329T C335T A341G |
| S3 | T318C |  |  |
| S4 | C383T |  |  |

\* Note, although a single nucleotide tide change is sufficient to support HRM genotyping, the Pan/99var15 virus contains multiple changes in a number of segments owing to use of alternative genotyping approaches during development of the IAV system.

To distinguish infected cells by flow cytometry, sequences encoding a 6xHIS or an HA epitope tag were added to the N-terminus of the IAV hemagglutinin protein, connected by a GGGGS linker sequence^23^. The 6xHIS tag was added to the Pan/99wt virus and the HA tag was added to the Pan/99var15 virus. The linker sequence provides flexibility so that the epitope tags do not interfere with HA protein folding. To ensure the tags were retained on the mature HA proteins, they were inserted after the signal sequence. It was not possible to add an epitope tag to reovirus, as this has been demonstrated in the literature and in our hands to cause growth defects (data not shown)^24^.

### Generation of virus stocks

Influenza A viruses were generated by reverse genetics from viral cDNA^25, 26^. Eight pPOL1 reverse genetics plasmids encoding the eight viral cDNAs were combined with four pCAGGS protein expression vectors encoding PB2, PB1, PA, and NP proteins. These plasmids were co-transfected into 293T cells using XtremeGene (Sigma-Aldrich) transfection reagent according to the manufacturer’s recommended procedure. At 16-24 h post transfection, the 293T cells were resuspended in growth medium and injected into the allantoic cavity of 9-11 day old embryonated chicken eggs (Hy-Line). Eggs were incubated 40 h in a humidified 33°C incubator. Incubation at 33°C was used because we have observed improved growth of the Pan/99 strain at this temperature, compared to 37°C ^27^. After chilling eggs overnight at 4°C, allantoic fluid was harvested, clarified of cell debris and aliquoted for storage at −80°C.

Mammalian orthoreoviruses were generated from viral cDNA cloned into the pT7 plasmid in BHK-T7 cells^28^. Plasmids containing each of the 10 viral cDNA’s and pCAG FAST P10 plasmid were transfected into BHK-T7 cells using TransIT-LT1 (Mirus)^29^. Cells were fed with an additional 500 μL SMEM on day 2. BHK-T7 cells were incubated at 37°C for a total of 5 days, after which virus was harvested with 3 freeze-thaw cycles at −80°C. Plaque assays were performed using viral lysates as previously described^30^. Reoviruses were amplified for two passages in L929 cells^30^. Purified virus stocks were made from second passage L929 lysates. Purification was performed as described previously using Vertrel XF extraction and a CsCl density gradient^31^. The band at 1.36 g/cm^3^ was collected and dialyzed exhaustively against virus storage buffer (15mM NaCl, 15mM MgCl_2_, 10mM Tris-HCl [pH 7.4]). The resultant purified material was stored in glass vials at 4°C for up to six months.

### Viral growth in guinea pigs

All animal experiments were approved by the Emory Institutional Animal Care and Use Committee under protocol number PROTO201700595, following the *Guide for the Care and Use of Laboratory Animals*^*32*^. Guinea pigs were inoculated intranasally with 10^3^ PFU of Pan99/wt or Pan99/var15 as described previously^33^. Nasal washes were collected on days 1, 2, 3, 4, and 5 post inoculation^33^. Virus titers in nasal washes were determined by plaque assay in MDCK cells.

### Primers

#### Reverse Transcription

For IAV, reverse transcription was performed using universal primers that anneal to the 3’ end of all eight IAV vRNA’s (GCGCGCAGC[A/G]AAAGCAGG) ^34^. Due to a lack of universally homologous sequences in reovirus, random hexamer primers (Thermo) were used in place of virus-specific primers.

#### High-resolution melt

Primers for quantitative PCR followed by high-resolution melt analysis were designed to flank the polymorphisms introduced into variant viruses. These primers were designed such that the amplicon size would be 65-110 nucleotides and annealing temperatures were 58-62°C. Primer sequences for IAV and reovirus are listed in Table 2. Primer mixes were made by combining 5 μL of the 100 μM forward primer stock and 5μL of the 100 μM reverse primer stock with 240 μL molecular biology grade water for a final concentration of 4 μM.

**Table 2:**
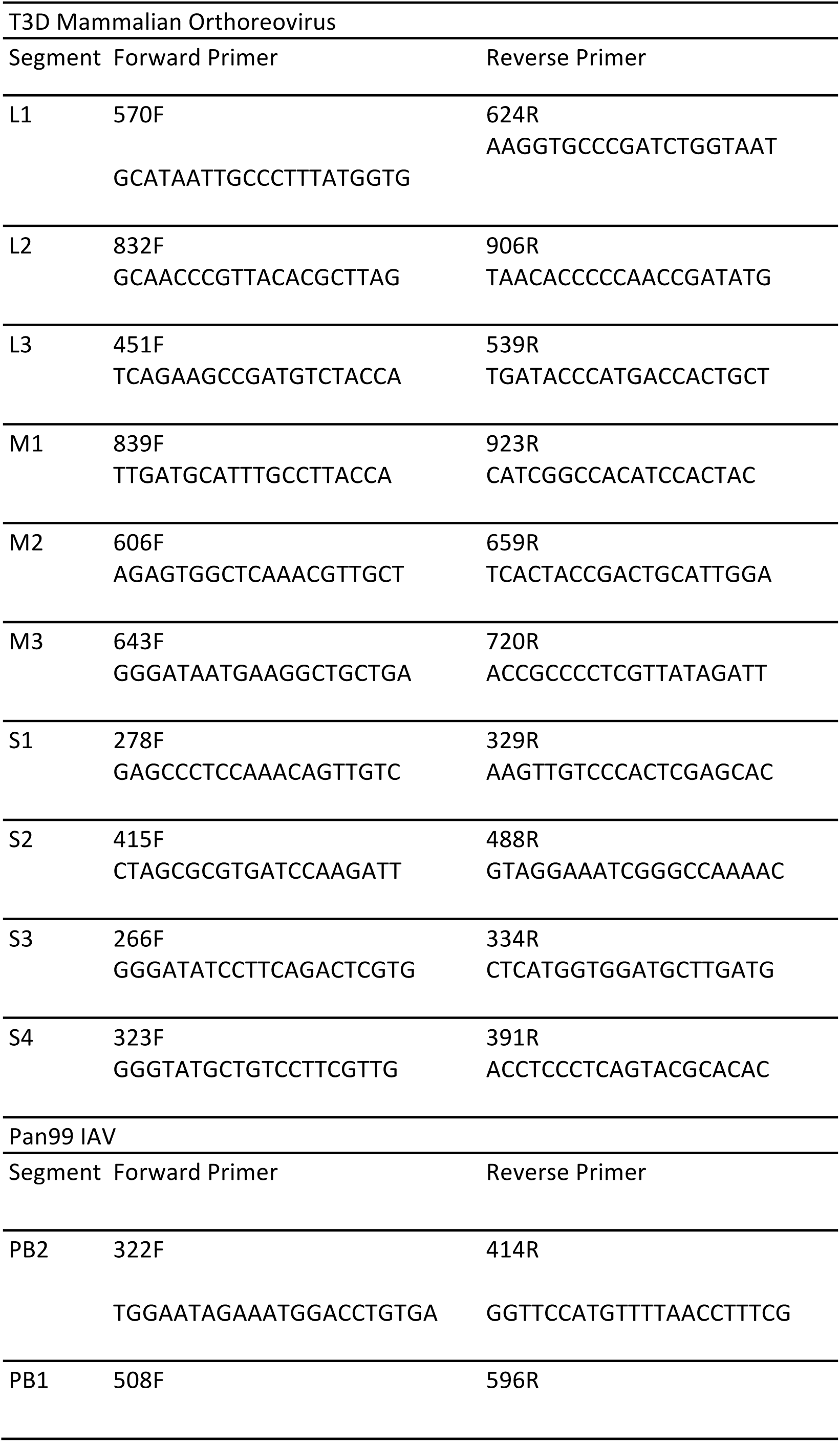
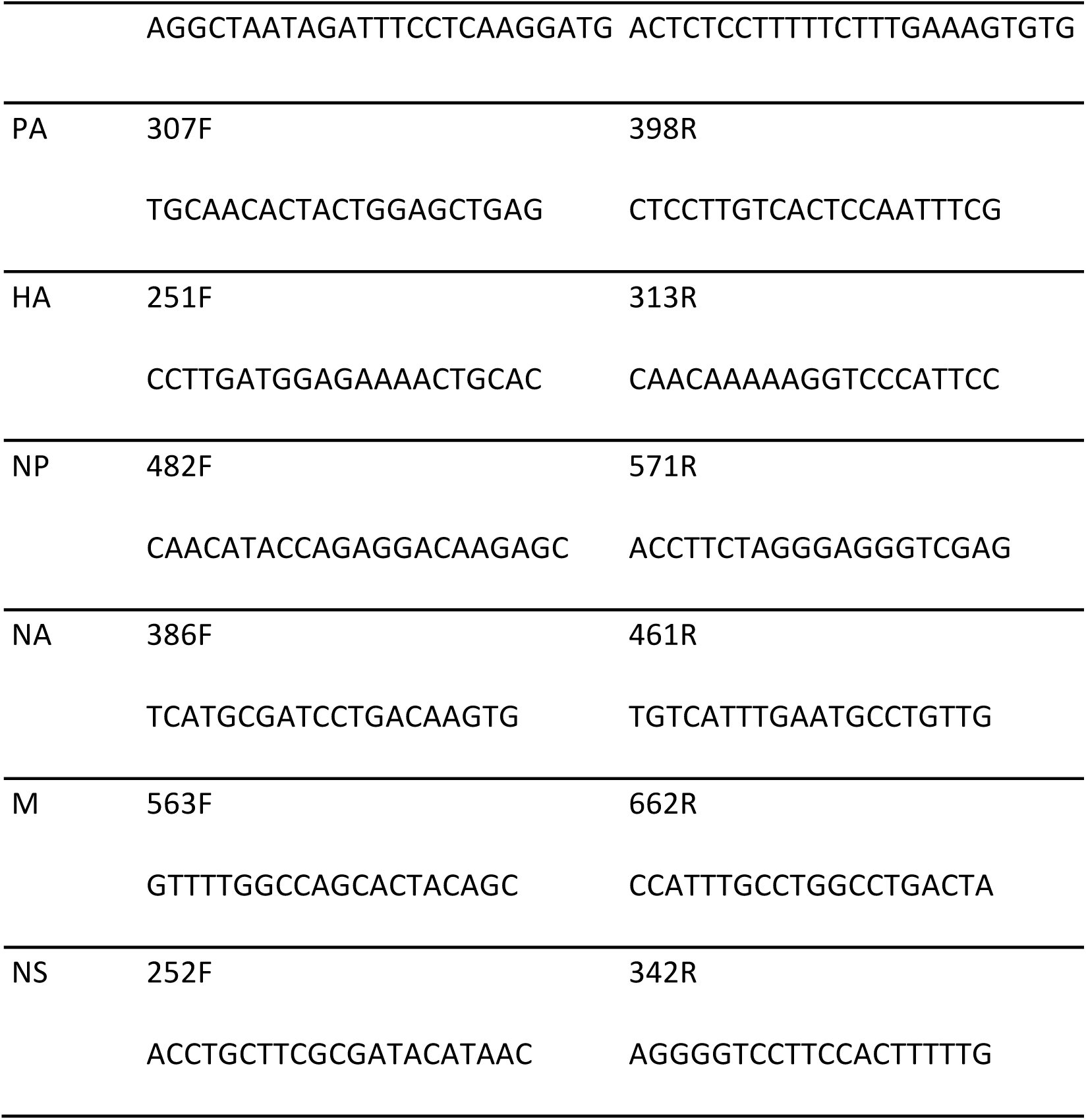
Primers used to generate amplicons for high-resolution melt analysis.

### Synchronized co-infection

#### IAV

Co-infections were performed with wild-type and variant (*wt/var*) viruses mixed at a 1:1 ratio and then diluted in PBS such that the total PFU/mL would give the desired multiplicity of infection. MDCK cells seeded at a density of 4×10^5^ cells/well in 6-well dishes 24 h prior were placed on ice, washed three times with 1X PBS, and inoculated with 200 μL of virus mixture in PBS. Dishes were moved to 4°C and rocked every 10 min for 45 min. After this attachment period, cells were put back on ice and washed 3 times with cold PBS. A 2 mL volume of warm virus medium (1X MEM with 100U/100 μg/mL penicillin/streptomycin, 0.3% bovine serum albumin [Sigma]) was added to each well, and plates were put into a 33°C incubator. At 3 h post infection (i.e. 3 h after warming), medium was removed and replaced with 2 mL warm virus medium supplemented with 20 mM NH_4_Cl and 50mM HEPES to prevent acidification of the endocytic compartment and thus prevent multiple cycles of infection. At 12 h post infection, supernatants were collected and stored at −80°C. Cells were harvested and prepared for analysis by flow cytometry.

#### Reovirus

Co-infections were similar to those for IAV, with a few modifications. L929 cells were seeded in 6-well dishes at a density of 5×10^5^ cells/well 24 h prior to infection. The virus inoculum was prepared in OPTI-MEM (Gibco), and cells were incubated in SMEM at 37°C. Rather than ammonium chloride, E64D protease inhibitor (Sigma) was added at a final concentration of 4 μM at 4 h post infection to block secondary infection. At 24 h post infection, three replicate wells of reovirus-infected cells at each MOI were harvested for flow cytometry, and the remaining three replicates were freeze-thawed 3 times at −80°C to release virus, and lysates stored at −80°C for future analysis.

### Flow cytometry to quantify infected cells

#### IAV

Cells were harvested by the addition of 200 μL trypsin (Corning) and, once cells were detached, 800 μL FACS buffer (1X PBS with 2% FBS). Cells were transferred to 1.5 mL tubes on ice, and pelleted by spinning at 1500 rpm for 3 minutes in a Beckman Coulter Microfuge 22R tabletop centrifuge. Supernatant was removed, and cells were washed two more times with 1 mL FACS buffer and 200 μL FACS buffer, respectively, pelleting and removing supernatant between washes. After washes, cells were resuspended in 50 μL stain buffer (FACS buffer containing Qiagen Penta-HIS Alexa Fluor 647 #35370 at a final concentration of 5 μg/mL and Sigma-Aldrich Monoclonal Anti-HA-FITC, Clone HA-7 #H7411 at a final concentration of 7 μg/mL) on ice in the dark for 35-45 minutes. Cells were washed twice with 200 μL of FACS buffer and resuspended in FACS buffer for analysis.

#### Reovirus

Cells were trypsinized and washed two times with FACS buffer, as above. Fixation, permeabilization, and staining were performed according to the BD Cytofix/Cytoperm protocol including a 15-minute block step with BD rat anti-mouse CD16/CD32 Fc block. To stain infected cells, a mouse monoclonal anti-σ3 antibody (clone 10C1) at a concentration of 1 μg/mL was added for 30 minutes at 4°C. After two washes, an AlexaFluor-647 conjugated donkey anti-mouse secondary antibody (Invitrogen) was added at a 1:1000 dilution.

In both virus systems, data was collected on a BD LSR II Flow cytometer running FacsDiva software. A minimum of 50,000 events was collected for each sample. Subsequent data analysis was performed using FlowJo (v10.1), gating for single cells. The threshold for positivity was determined based on a mock-infected control population stained with the relevant antibodies.

### Viral genotyping

#### Collection of clonal isolates

For both reovirus and IAV, samples stored at −80°C were thawed and plaque assays were performed as previously described^30^. Individual, well-isolated plaques were picked by aspirating the agar plug using a 1 mL pipette and deposited into 160 μL PBS in a 96-well assay block with 1 ml capacity wells (Costar 3958). From each sample, 21 plaques were picked for IAV, while 32 were picked for reovirus. Additionally, a single wild-type and a single variant plaque were included as controls for each series of 21 or 32 plaque picks. Assay blocks containing plaque isolates can be sealed and stored at −20°C or used directly for RNA extraction.

#### RNA extraction

RNA was extracted directly from agar plugs. Frozen assay blocks were thawed in a 37°C water bath and spun down at 2000 rpm for 2 min in a Heraeus Megafuge 16 tabletop centrifuge equipped with Thermo M-20 swinging bucket plate rotor. The Zymo *Quick*-RNA Viral Kit extraction protocol was followed using 96-well plates. Filter and collection plates were provided with the kit. No DNA/RNA Shield was used. Samples were eluted in 40 μL nuclease-free water into MicroAmp Optical 96-well reaction plates (Applied Biosystems). RNA can be covered and stored at −80°C or used directly for reverse transcription.

#### Reverse Transcription

Working on ice, a 12.8 μL volume of each viral RNA sample was combined with Maxima RT buffer at a final concentration of 1X, dNTP’s at a final concentration of 0.5 mM, either IAV primer at a final concentration of 0.3 μM or random hexamers (Thermo SO142) at a final concentration of 5 μM for reovirus, 100 U Maxima RT (Thermo), and 28 U RiboLock RNAse inhibitor (Thermo). Total reaction volume was 20 μl. Samples were capped, mixed by vortexing, and spun down briefly. Reactions were incubated at 55°C for 30 minutes and 85°C for 10 minutes in a BioRad T100 thermocycler. cDNA can then be stored at −20°C or used directly for qPCR and high-resolution melt analysis.

#### qPCR and high-resolution melt analysis

Viral cDNA was used as a template in qPCR reactions. Separate reactions were set up with primers targeting each of the viral gene segments. First, master mixes were made by combining appropriate primer mixes (see Table 2) with BioRad Precision Melt Supermix at volumes sufficient for the number of samples plus 15% extra. For each well, 0.5 μL of a 4 μM primer mixture containing both the forward and reverse primers was added to 2.5 μL Supermix. A 3 μl volume of this master mix was loaded into a 384 well plate (BioRad HSP3805) using a multichannel pipette according to the layouts in Figure 2. A 2 μl volume of cDNA diluted 1:4 (IAV) or 1:5 (reovirus) in molecular biology grade water (Invitrogen) was added to the 384 well plate. Plates were centrifuged at 2600 rpm in a Heraeus Megafuge 16 tabletop centrifuge equipped with Thermo M-20 swinging bucket plate rotor for 3 minutes to collect liquid in the bottom of wells and remove bubbles. qPCR and melt analysis were performed using a BioRad CFX384 Real-Time PCR Detection System. Amplicons were generated by initial denaturation at 95°C for 2 min, then 40 cycles of 95°C for 10 s and 60°C for 30 s. Melting properties of PCR amplicons were examined by heating from 67°C to 90°C in 0.2°C increments. Successful amplification of targets was verified in CFX Manager software (BioRad). Melt curves were analyzed using Precision Melt Analysis software (BioRad) to determine viral genotypes.

**Figure 1.**
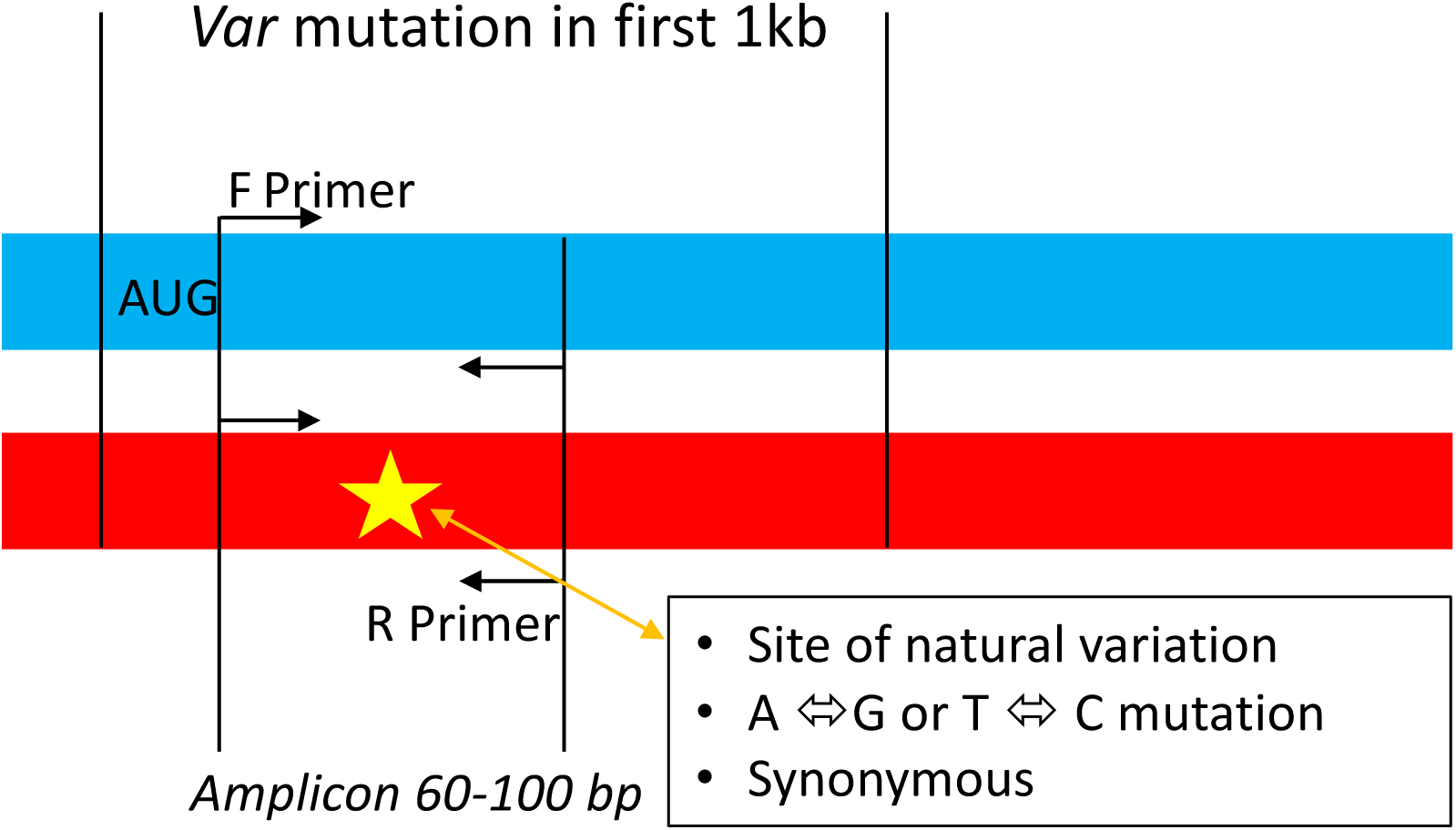
Design of variant mutations. Genome segments are indicated by blue (wt) and red (var) bars. The single nucleotide polymorphism is indicated by a star in the variant segment. Vertical lines indicate the 1 kb region (beginning at the start codon) in which the polymorphism was introduced, and the borders of the amplicon used in qPCR and subsequent high resolution melt analysis. Arrows indicate the directionality of the primers used during qPCR. Criteria used in the selection of the variant mutation position are indicated in the box on the lower right.

**Figure 2.**
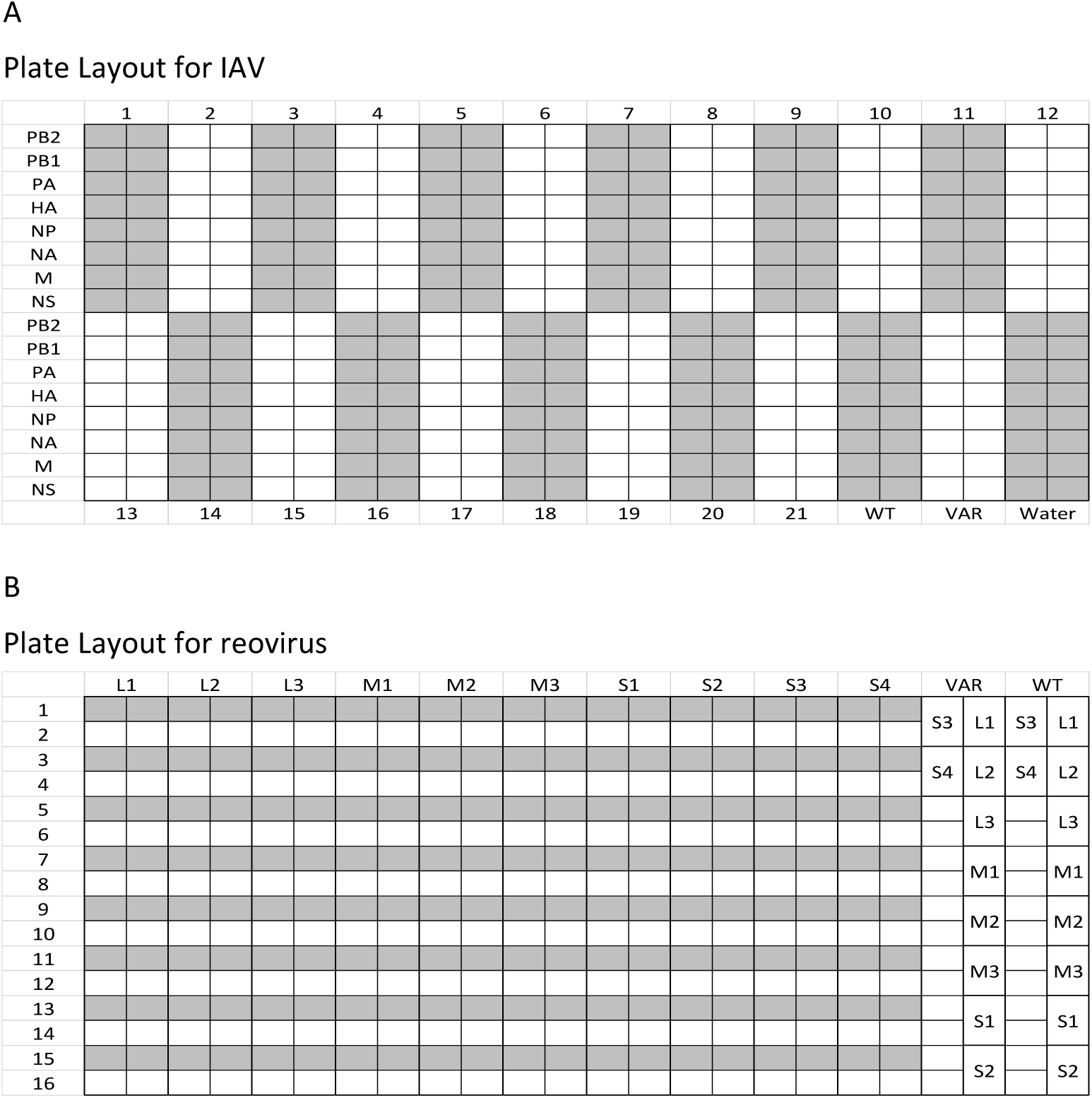
Example 384-well plate layouts for high-resolution melt analysis. A) Example plate layout for the analysis of IAV reassortment. Each plate holds 21 unknown samples (indicated by numbers at the top and bottom of each schematic plate), wt and var positive controls (bottom right) and a negative control in which water was loaded in place of cDNA (bottom right). Each of the 8 segments are analyzed in duplicate for each sample, in rows as indicated at the left with segment designations. B) Example plate layout for the analysis of reovirus reassortment. Each plate holds 16 samples (indicated by numbers to the left), and wt and var positive controls (right side of the diagram). Each of the 10 segments are analyzed in duplicated, indicated by segment identification across the top and in wells for positive controls. Since 32 isolates are analyzed for reovirus reassortment, two plates must be used.

## Results

### Marker mutations in variant viruses do not cause fitness defects

To quantify reassortment in the absence of selection bias, it is important to ensure that the parent viruses do not differ in fitness. If one virus is fitter, that parental genotype and segments from it are likely to predominate in the progeny virus population due to uneven amplification. This outcome may be of interest in some contexts, but obscures quantitative analysis of reassortment itself. For IAV, fitness of wild type and variant viruses was assessed in a guinea pig model. Groups of four guinea pigs were inoculated intranasally with Pan/99wt or Pan/99var15 virus and nasal washes were performed daily to monitor viral load. The two viruses showed very similar growth, indicating minimal phenotypic impact of the changes introduced into the viral genomes (Figure 3). For mammalian orthoreovirus, single-cycle growth in L929 cells was evaluated to compare the fitness of the variant and wild type viruses. Again in this system, the variant strain was not attenuated relative to the wild-type counterpart (Figure 3). The highly homologous genotypes and comparable fitness of parental viruses are expected to result in similar fitness of reassortant progeny, allowing for an unbiased assessment of reassortment frequency.

**Figure 3.**
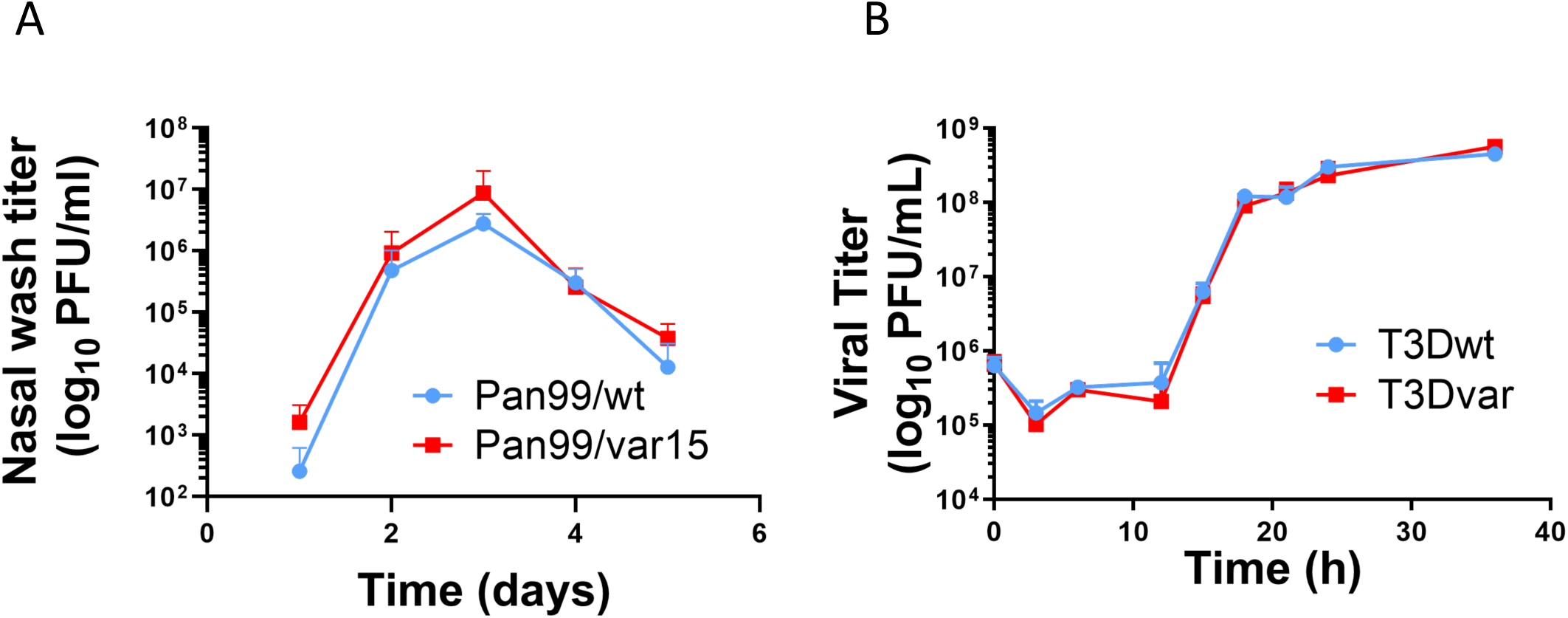
Single nucleotide changes do not detectably alter variant virus growth. (A) Pan/99 IAV wild-type and variant multi-cycle growth were analyzed in guinea pigs (N=4 Pan/99wt, N=3 Pan/99var15) over the course of 5 days. (B) T3D mammalian orthoreovirus wild-type and variant virus growth were analyzed under single-cycle conditions in L929 cells over the course of 36 h (N=3 for both viruses). Error bars represent standard deviation.

### Flow cytometry allows quantification of the proportion of cells infected

Owing to the presence of non-plaque forming particles in virus populations and routine experimental error, calculation of MOI based on plaque forming units does not always allow an accurate estimation of the proportion of a cell population that is infected. Flow cytometry analysis targeting viral antigens allows a more direct method to monitor infection levels.

In the case of IAV, the addition of different epitope tags to wt and var HA proteins allowed quantification of single- and co-infected cells within the population. Four distinct populations were observed representing uninfected, wild-type virus infected (expressing 6x HIS tag), variant virus infected (expressing HA tag), or wild type plus variant co-infected cells (Figure 4). Assessment of the proportion of cells in a population that are co-infected is useful, as co-infected cells are the only ones capable of producing reassortant progeny.

**Figure 4.**
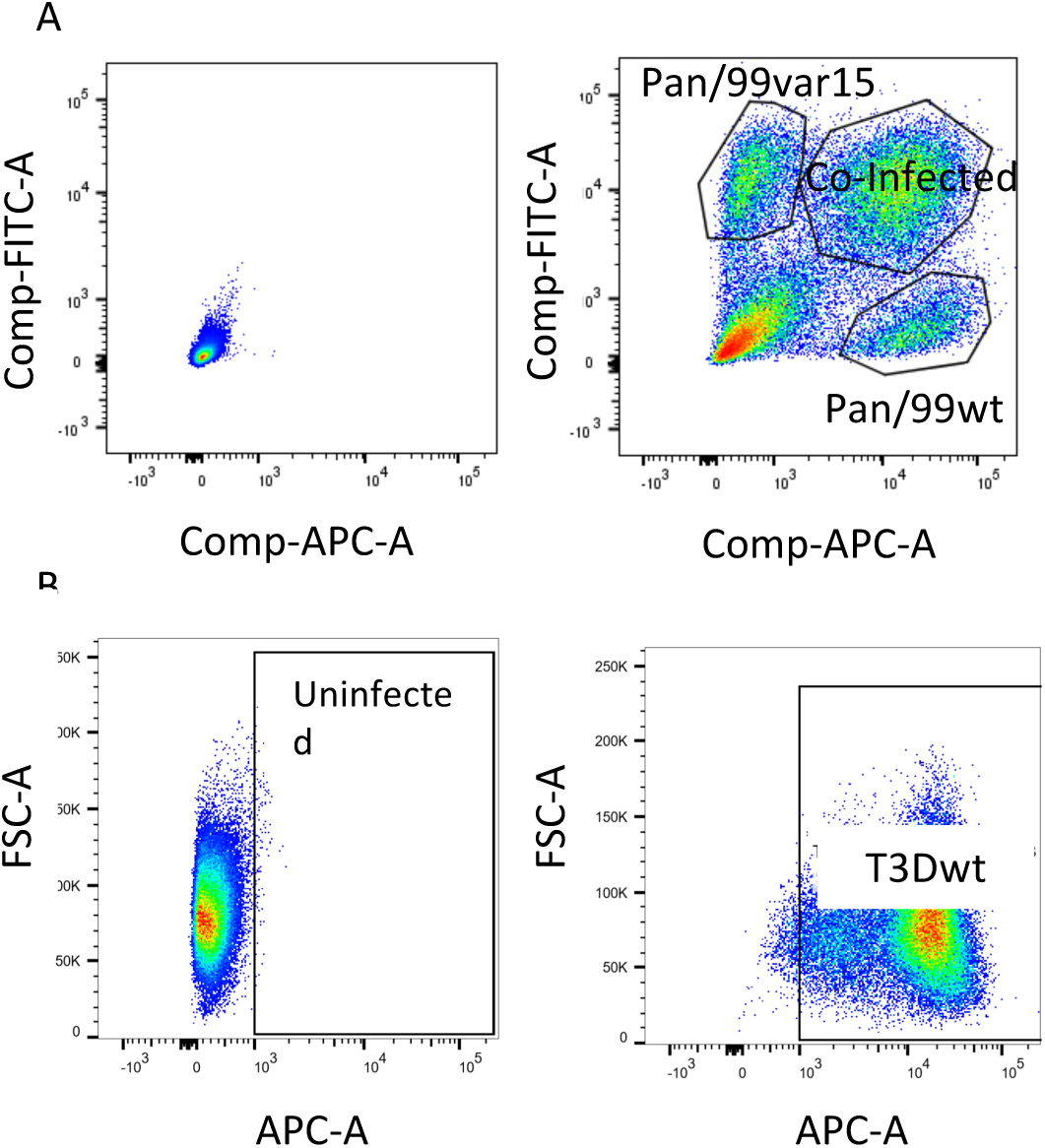
Flow cytometry allows for the quantification of infected and co-infected cells in a population. (A) Uninfected cells (left) were used to determine the appropriate gate locations for IAV infected cells (right). Populations of cells infected by a single virus are indicated by the top left- and bottom rightmost gates. Co-infected cells expressing both the HA and 6xHIS epitope tags are shown in the top rightmost gate. (B) Uninfected cells (left) were used to determine the appropriate gate location for reovirus infected cells (right). The population shift indicates 98.6% infected cells divided between two populations, representing high and low levels of viral antigen expression.

Since insertion of epitope tags was not feasible in the reovirus system, flow cytometry of reovirus-infected cells was used to evaluate the proportion of cells that were infected, but no direct measurement of co-infected cells was made. Infected cells were detected using a primary antibody against viral structural protein σ3 (10C1), and compared to an uninfected, stained control. Infected cells cluster within two groups with low and high antigen expression, respectively (Figure 4). To estimate the percentage of infected cells that are co-infected, Poisson statistics can be used, assuming an equal proportion of wild-type and variant viruses were present in the initial infection.

### High-resolution melt analysis allows determination of viral genotypes

High-resolution melt analysis allows detection of the nucleotide changes that differentiate wt and var viruses, and can therefore be used for rapid assignment of wt or var genotypes to all segments present in clonal virus isolates. To this end, qPCR was performed with the cDNA of each viral isolate split into eight (IAV) or ten (reovirus) separate duplicate reactions, each containing primers targeting a different segment. Following qPCR, samples with C_t_ values below 35 were used for melt analysis. The included wt and var controls were used as references and clusters based on similarity of T_m_ and melt curve shape were generated within BioRad High Precision Melt software (Figure 5). The distinct melt curves of wt and var amplicons indicate that the silent mutations introduced were sufficient for identification of the parental origins of each segment (Figure 5). In practice, we find that a minimum T_m_ difference of 0.15°C is needed to consistently differentiate between wt and var amplicons. Applying this approach to each segment in turn allows the full genotype of each clonal plaque pick to be determined.

**Figure 5.**
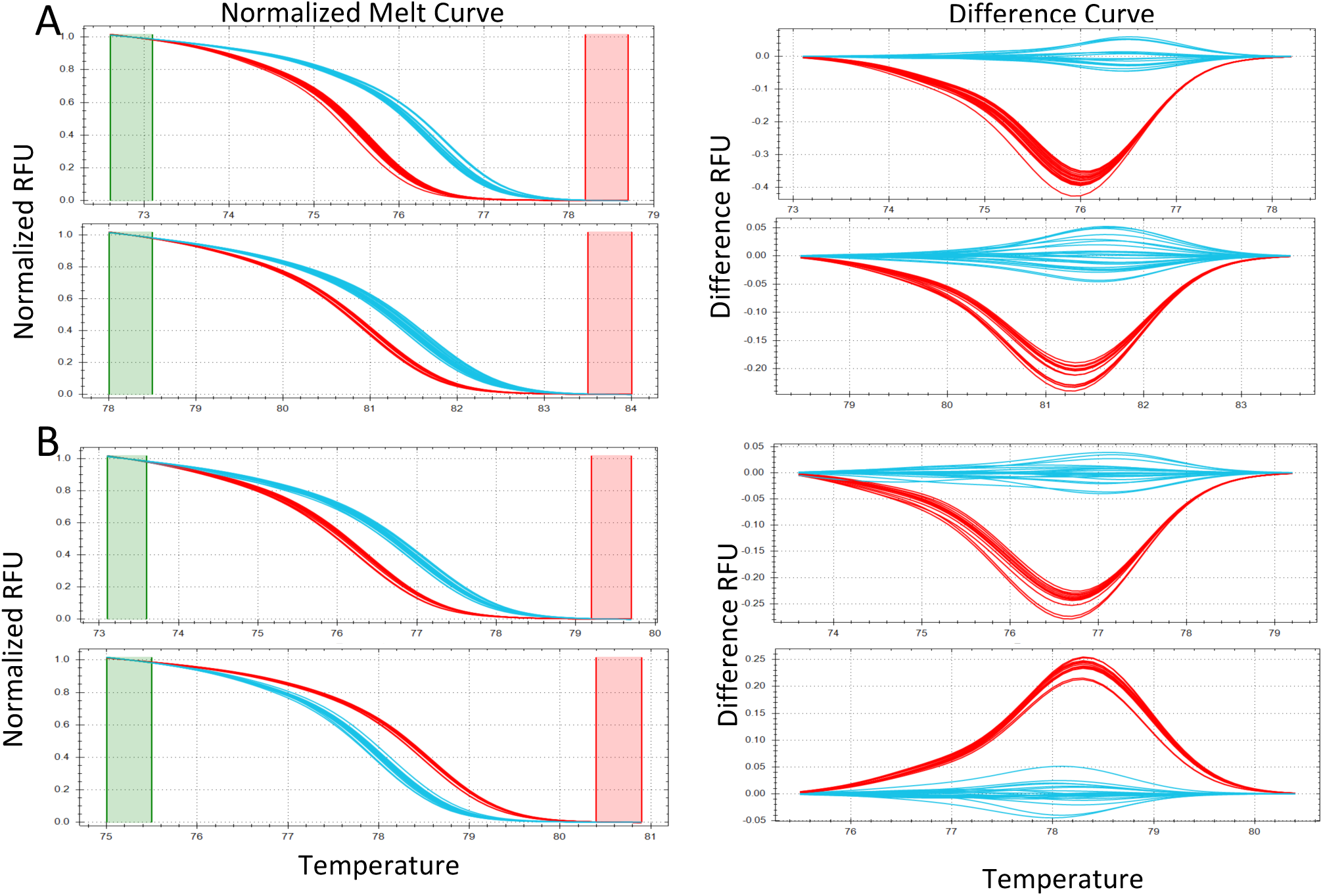
Melt curves allow the determination of parental segment origin. Melt curves on the left indicating relative fluorescent units (RFU) as a function of temperature were used to generate the difference curves on the right, which enabled differentiation of wt and var segments in the clusters. Wild-type (blue) and variant (red) controls were used to determine the parental origins of each segment. Melt curves of two representative segments from IAV (A) and reovirus (B) are shown.

Full genotypes are depicted in reassortment tables where each column is a separate genome segment, and rows represent clonal isolates (Figure 6). Occasionally, high resolution melt analysis yielded unclear results, with a given amplicon clustering neither with wt nor var controls. Such results were recorded as indeterminate (white boxes in Figure 6). Viral isolates were excluded from analysis if there were more than two segments omitted due to unclear melt curves. If more than 20% of replicates were omitted from analysis, data collection was repeated starting from plaque picks.

**Figure 6.**
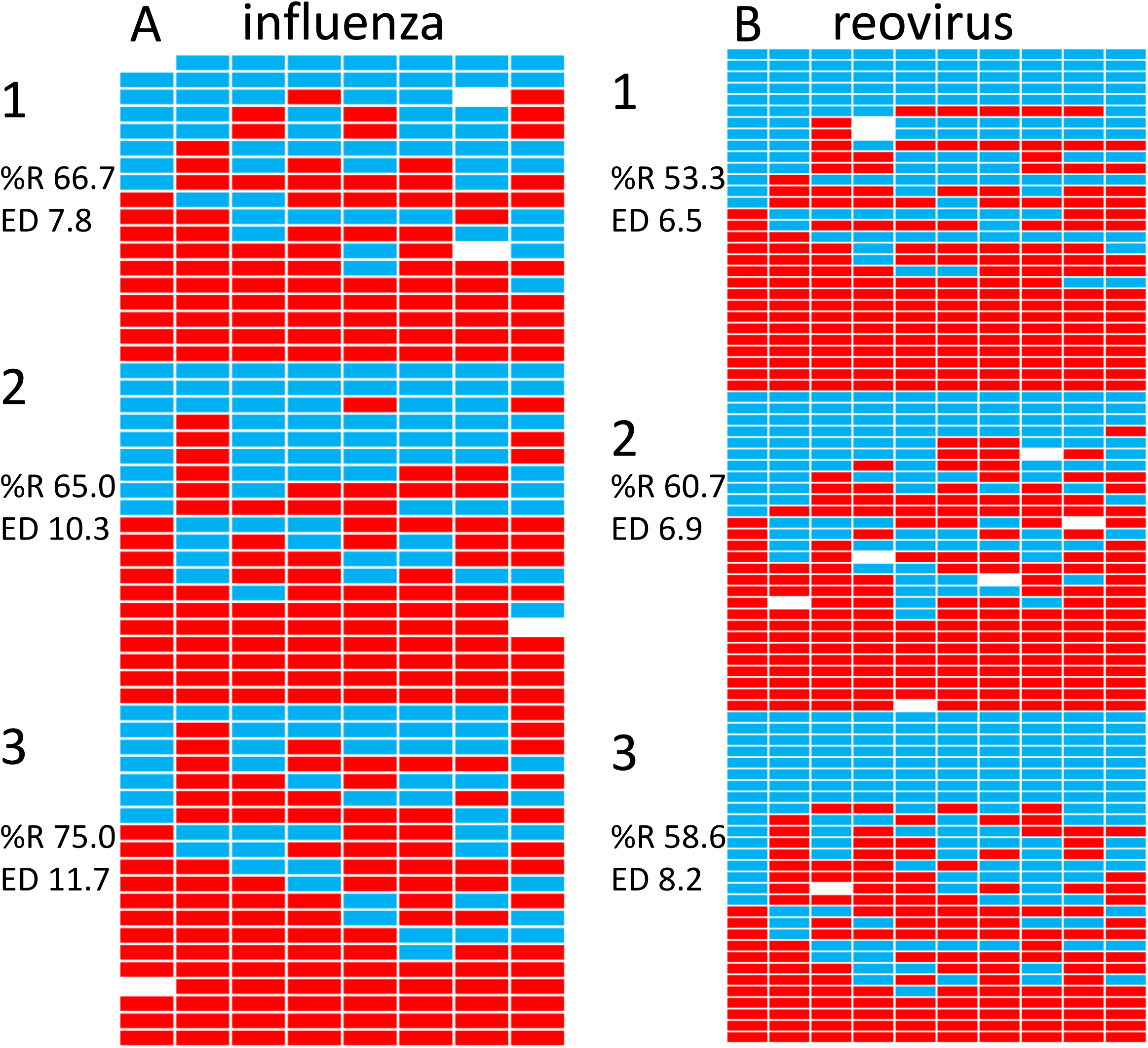
Reassortment tables provide visual representation of progeny virus genotypes. Each co-infection was performed in triplicate, and 21 (IAV) or 32 (reovirus) plaques were genotyped from each replicate. Blocks of genotypes numbered 1, 2 and 3 represent each replicate co-infection. Each row corresponds to a separate plaque isolate, and each column represents a gene segment (IAV segment order: PB2, PB1, PA, HA, NP, NA, M, NS; Reovirus segment order: L1, L2, L3, M1, M2, M3, S1, S2, S3, S4). Representative data from co-infections performed at a single MOI of both IAV (A) and reovirus (B) are shown. An MOI of 0.6 is shown for IAV, and an MOI of 3.16 is shown for reovirus. The calculated percent reassortment (%R) and the diversity (ED) as measured by Simpson’s index are indicated for each replicate.

Following assembly of genotype tables, the percent reassortment observed in each sample is calculated as 100 times the number of reassortant clones identified divided by the total number of clones genotyped. To evaluate the relationship between infection and reassortment, the calculated percent reassortment can be visualized as a function of the percent infected cells as calculated by flow cytometry (Figure 7). For this quantitative assessment of reassortment to be meaningful, it is important that co-infections be performed under single cycle conditions. As noted in the Materials and Methods section, this was achieved for IAV using addition of ammonium chloride to cell culture medium at 3 h post-infection, and for reovirus using E64D protease inhibitor added at 4 h post-infection. Blocking secondary spread of progeny virus ensures that detected frequencies of reassortant viruses reflect the efficiency of reassortment, rather than the efficiency of amplification. In contexts where infection cannot be limited to a single cycle, such as in vivo, analysis of genotypic diversity (as described below) is more appropriate than a simple readout of percent reassortment.

**Figure 7.**
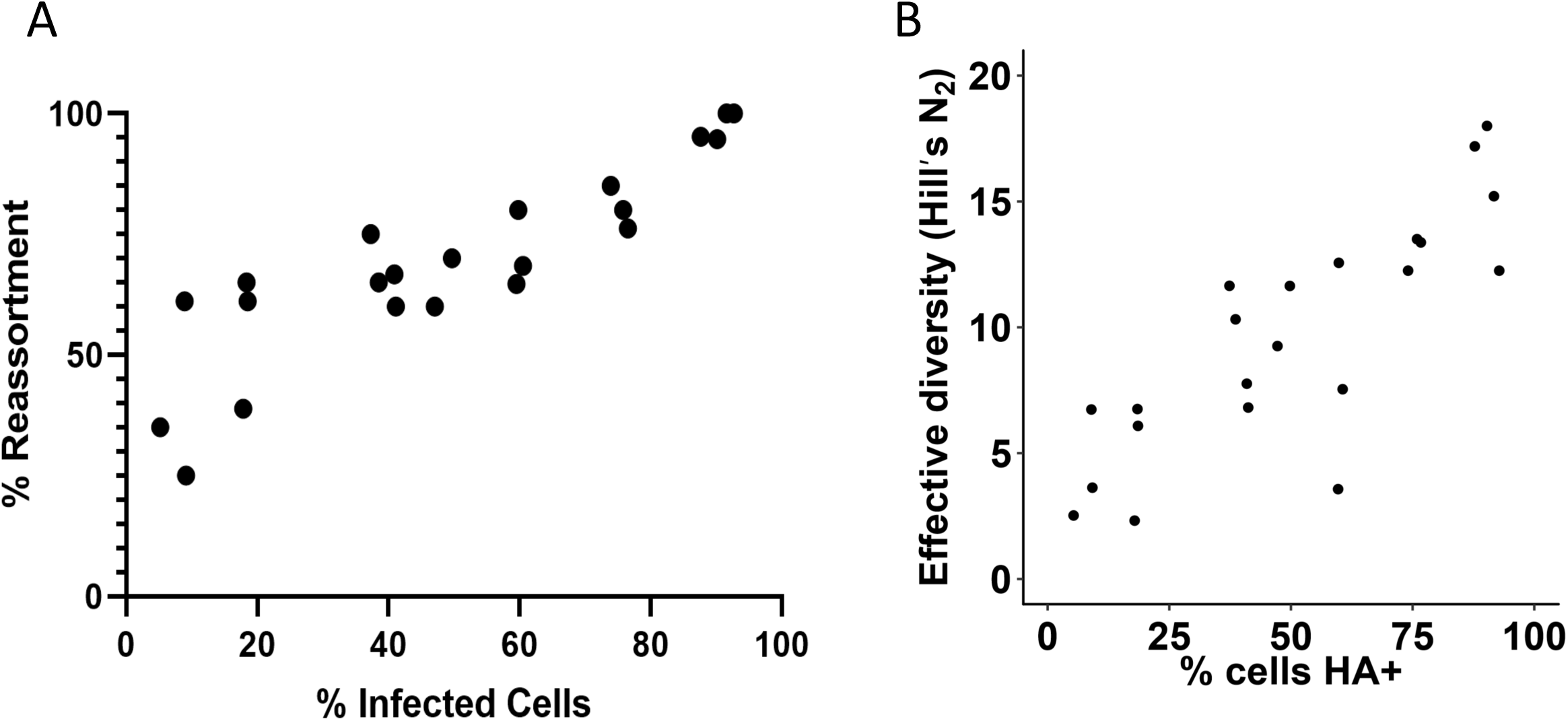
Comparison of infection levels and reassortment. Quantification of infected cells by flow cytometry can be combined with reassortment quantification to give insight into the dependence of reassortment on effective viral dose. (A) Percent reassortment as a function of the percentage of cells in the population infected with Pan/99 IAV, as detected by HA expression in flow cytometry. (B) The effective diversity of Pan/99 isolates after co-infections in A549 cells was determined. Diversity is shown as a function of % infected cells, expressed as % HA-positive (expressing HA-labelled HA protein, 6xHIS-labelled HA protein, or both). Diversity increases more than five-fold from the lowest % infection to the highest.

### Diversity analysis quantifies richness and evenness of reassortant population

A sample in which a single reassortant genotype is detected repeatedly would have high percent reassortment despite having low genotypic diversity. When using a wt/var co-infection system, this situation is unlikely to arise due to selection, but can nevertheless occur under conditions where stochastic effects are strong (e.g. owing to within host bottlenecks in vivo). Here, the percent reassortment readout is not highly relevant and a more sophisticated analysis of genotypic diversity is needed.

To quantify the diversity of genotypes present, Simpson’s index (given by D = sum(p_i_^2^), where p_i_ is the proportional abundance of each genotype) was used (Figure 7). This approach accounts for both the raw number of species (richness) and variation in the abundance of each (evenness), and is sensitive to the abundance of dominant species. To determine effective diversity, the Simpson index value of each sample was converted to a corresponding Hill number, N_2_ = 1/D. The Hill number N_2_ is equivalent to the number of equally abundant species needed to generate the observed diversity in a sample community, and is particularly useful because it scales linearly (i.e., a virus population with N_2_ = 10 species is twice as diverse as one with N_2_ = 5). Hill’s N_2_ therefore allows a more intuitive comparison between populations and is suitable for statistical analysis by basic linear regression methods^35^.

## Discussion

Here we outline a conceptually simple approach to accurately quantify reassortment between co-infecting segmented viruses in the absence of selection bias. Our strategy utilizes reverse genetics derived parental viruses designed to allow both unbiased reassortment and streamlined genotyping. This approach overcomes limitations of previous methods in which quantitative analysis of reassortment was impeded by fitness differences among progeny viruses. The genotyping technology employed furthermore improves upon more cumbersome procedures involving gel electrophoresis or temperature-sensitive mutants. This method is useful for fundamental studies of reassortment and other interactions within virus populations.

High resolution melt analysis can also be applied to viral genotyping in systems where highly divergent parental viruses are allowed to reassort. Although fitness differences among progeny viruses will obscure the quantitative assessment of reassortment in such an experiment, monitoring the combined outcomes of reassortment and selection is often highly relevant for assessing the public health risks posed by reassortment of parental strains of interest^22, 36-38^. Highly divergent parental sequences are likely to exhibit detectable differences in melting properties, conducive to HRM analysis. It is important to note, however, that reciprocal changes within the region targeted for amplification and melt analysis will nullify differences in melting properties and therefore prohibit detection. Short regions with fewer single nucleotide polymorphisms are less likely to contain reciprocal changes, and should be selected for amplification. In addition, regions of high homology must border the target region, as the same HRM primer set must be used for each parent. In practice for IAV, when considering reassortment between strains of differing subtypes, we found that HRM can be successfully applied to genotype the six non-HA, non-NA segments. The low sequence identity across HA and NA subtypes precluded common primer design and an alternative genotyping approach was used for these segments^22^.

Whole genome sequencing of clonal viral isolates is an alternative to the HRM approach that has been used recently to identify reassortant viruses^39^. Next generation sequencing (NGS) allows parallel sequencing of all viral gene segments, which greatly reduces effort compared to classical Sanger sequencing. In addition, because viral genomes are typically small, the costs of NGS can be reduced by combining the barcoded cDNA derived from multiple isolates into a single sequencing lane. NGS can be applied to any pairing of parental viruses and may be more feasible than HRM where parental viruses are highly divergent and identical HRM primers cannot be generated for all segments. However, in experiments using matched parental viruses, such as the wt/var system, the HRM approach simplifies data collection. Whole genome sequencing requires additional preparation steps including the pre-amplification of cDNA, fragmentation, and library generation. Additionally, customized bioinformatics approaches are necessary for NGS data analysis, but not required in the HRM approach.

Segmented viruses utilize a variety of replication strategies which may impact their potential to undergo reassortment. For this reason, quantification of reassortment not only informs studies of viral diversification and evolution, but can also offer insight into fundamental aspects of the virus life cycle. For example, members of the *Cystoviridae* such as phi6 bacteriophage use genome packaging as an integral part of the assembly mechanism^40^. Successful assembly can only occur if a full complement of three gene segments is incorporated into a virion. In co-infections between highly divergent parents, an inability to package any of the three segments results in attenuation, which has the potential to limit emergence of reassortant viruses. Conversely, only ∼1% of eight-segmented influenza A viruses replicate a full complement of segments within the infected cell, so co-infection is often necessary for successful completion of the viral life cycle^41^. As co-infected cells are the only ones capable of yielding reassortant progeny, a requirement for co-infection increases the likelihood that reassortment will occur. Progeny resulting from the co-infection of a cell with two viruses bearing incomplete genomes are necessarily reassortant. The rigidity of packaging mechanisms and the prevalence of incomplete genomes in other virus families, such as the *Reoviridae*, remains incompletely understood, as does the frequency of reassortment in these systems. Utilization of the strategy outlined herein for monitoring reassortment may open up further avenues of study with respect to segmented virus replication mechanisms and population dynamics.

## Acknowledgements

This work was funded in part by the NIH/NIAID Centers of Excellence for Influenza Research and Surveillance (CEIRS) contract HHSN272201400004C and R01 AI 125258. We thank Bernardo Mainou and Nathan Jacobs for helpful discussion.

